# An ancestral genomic sequence that serves as a nucleation site for *de novo* gene birth

**DOI:** 10.1101/2022.01.12.475983

**Authors:** Nicholas Delihas

## Abstract

A short non-coding sequence present between the gamma-glutamyltransferase 1 (*GGT1*) and gamma-glutamyltransferase 5 (*GGT5*) genes, termed a spacer sequence has been detected in the genomes of *Mus musculus*, the house mouse and in *Philippine tarsier*, a primitive ancestral primate. It is highly conserved during primate evolution with certain sequences being totally invariant from mouse to humans. Evidence is presented to show this intergenic sequence serves as a nucleation site for the initiation of diverse genes. We also outline the birth of the human lincRNA gene *BCRP3* (BCR activator of RhoGEF and GTPase 3 pseudogene) during primate evolution. The gene developmental process involves sequence initiation, addition of a complex of tandem transposable elements and addition of a segment of another gene. The sequence, initially formed in the Old World Monkeys such as the Rhesus monkey (*Macaca mulatta*) and the baboon (*Papio anubis*), develops into different primate genes before evolving into the human *BCRP3* gene; it appears to also include trial and error during sequence/gene formation. The protein gene, *GGT5* may have also formed by spacer sequence initiation in an ancient ancestor such as zebrafish, but spacer and *GGT5* gene sequence drift during evolution produced a divergence that precludes further assessment.

**Author summary:** For a number of decades researchers have been interested in how genes evolve and a number of mechanisms of gene formation have been defined. This manuscript describes a different process of gene formation, that of a small DNA sequence that does not code for a gene but serves as a nucleation site for the initiation of *de novo* gene formation. This non-coding DNA sequence appears to have been in existence for about hundred million years or more and has formed the basis for the birth of diverse genes during evolution of the primates. The questions of how and why new genes are born are important in terms of revealing how organisms, especially primates, progress to greater complexity during evolution; the question of “how” is particularly relevant to the creation of biological information *ab initio* during prebiotic and early cellular evolution.

## Introduction

Protein genes are created by varied processes that include gene duplication [1–5], retrogenes [6] and *de novo* formation [6–12]. With respect to the latter, Knowles and McLysaght [8] first reported that several human protein-coding genes arose by a *de novo* mechanism, and Wu et al [9] identified 60 protein-coding genes that are also born by a *de novo* process. Less has been reported on origins of long intergenic noncoding RNA (lincRNA) genes. However, some examples are lincRNA genes created from pseudogenized protein genes [13] and lincRNA family genes formed by gene duplication [14]. In addition, the formation of a new human lincRNA gene by transcriptional readthrough has been reported. The work of Rubino et al [15] shows that by use of the transcriptional apparatus of an existing gene and transcriptional readthrough to a small intergenic sequence that represents a functional unit, a new gene is created. This new gene is thought to participate in regulation of the immune system. This study has similarities to the work of Shiao et al [16] concerning *de novo* acquired 3’ UTRs that may play important functions of retrogenes, and that of Stewart and Rogers [17] in terms of the recruitment of non-coding sequences with chromosomal rearrangements and the resultant formation of new protein genes. Thus far, the creation of lincRNA genes appears similar to that of protein genes.

Here we describe a different process of *de novo* gene birth. It was previously thought that the lincRNA *FAM247* family gene sequence may serve as a nucleation site for new gene birth [18]. However, outlined here is a non-coding DNA sequence, termed a spacer sequence that is situated between the gamma-glutamyltransferase 1 (*GGT1*) and gamma-glutamyltransferase 5 (*GGT5*) genes. It is present in the rodent house mouse (*Mus musculus*) and ancestral prosimian primitive primates such as *Philippine tarsier* (*Carlito syrichta*) and is evolutionarily conserved in the genomes of all higher primates. It consists of less than 4000 bp, and in many species can contain small sections of the FAM247 sequence. We show that this spacer sequence is a nucleation site for new gene formations. We find that the 3’ ends of spacers are sites for the in initiation of *de novo* sequence growth with the creation of diverse genes during primate evolution. In addition, the chimpanzee genome provides an example of the combination of spacer sequence duplication and *de novo* gene formation at the duplicated genomic locus, which is partly analogous to chromosomal rearrangements and the resultant generation of *de novo* genes described by Rogers and Stewart [17]. Eight experimentally and/or computationally determined genes have been detected that stem from spacer sequences during primate evolution and all sequences starts with the elongation of the FAM247 sequence.

Also presented here are the evolutionary formations of two human long non-coding RNA genes, the lincRNA gene *BCRP3* (the BCR pseudogene 3), and the *FAM247A,C,D*, long intergenic RNA family genes, and propose a model for the formation of the BCRP3 sequence in the Rhesus monkey. With these genes, a trial and error process to produce the complete sequence appears to have occurred in several ancestral primates. We also discuss the presence of a significant length of conserved transposable elements (TEs), Alu/LINE TE tandem repeats found in the BCRP3 sequence. These tandem repeats pose interesting questions of origin and function. Aside from non-coding RNA genes, it is possible that the *GGT5* protein gene, whose sequence also begins with an FAM247 sequence and is found in non-mammalian ancestors, may also have formed via spacer sequence initiation. The zebrafish genome may be an ancestral example where *GGT5* was born, but the spacer sequence significantly diverged during evolution, which makes further assessment of spacer involvement in *GGT5* formation difficult.

## Results

### Properties of spacer sequences

Computational alignment and search programs were used to analyze genomes of primates and other species. The primitive early primate, *Philippine tarsier* genome was found to display a small genomic spacer sequence (2872 bp) situated between the protein genes *GGT1* and *GGT5* (Fig. 1). The spacer sequence between *GGT1* and *GGT5* in the house mouse *Mus musculus* genome is also shown. A large expansion at this genomic region occurred during primate evolution as the Rhesus monkey sequence between genes *GGT1* and *GGT5* shows an increase in size to 216,200 bp; this sequence expansion is on chr10 and the sequence is also found inverted with chromosomal rearrangements (Fig. 1). The chimpanzee genome continued this genomic expansion with an increase to 343,330 bp; the human genome maintained most of this sequence but decreased by ~10%. The spacer sequence between *GGT1* and *GGT5* of the primitive primate *Philippine tarsier* is found in the higher primates linked to the *GGT1* gene after genomic expansion. The expanded genomic regions contain duplicated sequences that have provided for the formation of a number of new genes or family of genes. However, of significance, the *GGT1*-spacer sequence 3’ ends are found to be focal points, or nucleation sites where diverse genes and/or sequences originate from the *GGT1*-spacers in various primate species, the spacer 3’ end serving as the starting point for growth of new sequences and/or genes.

**Fig. 1.**
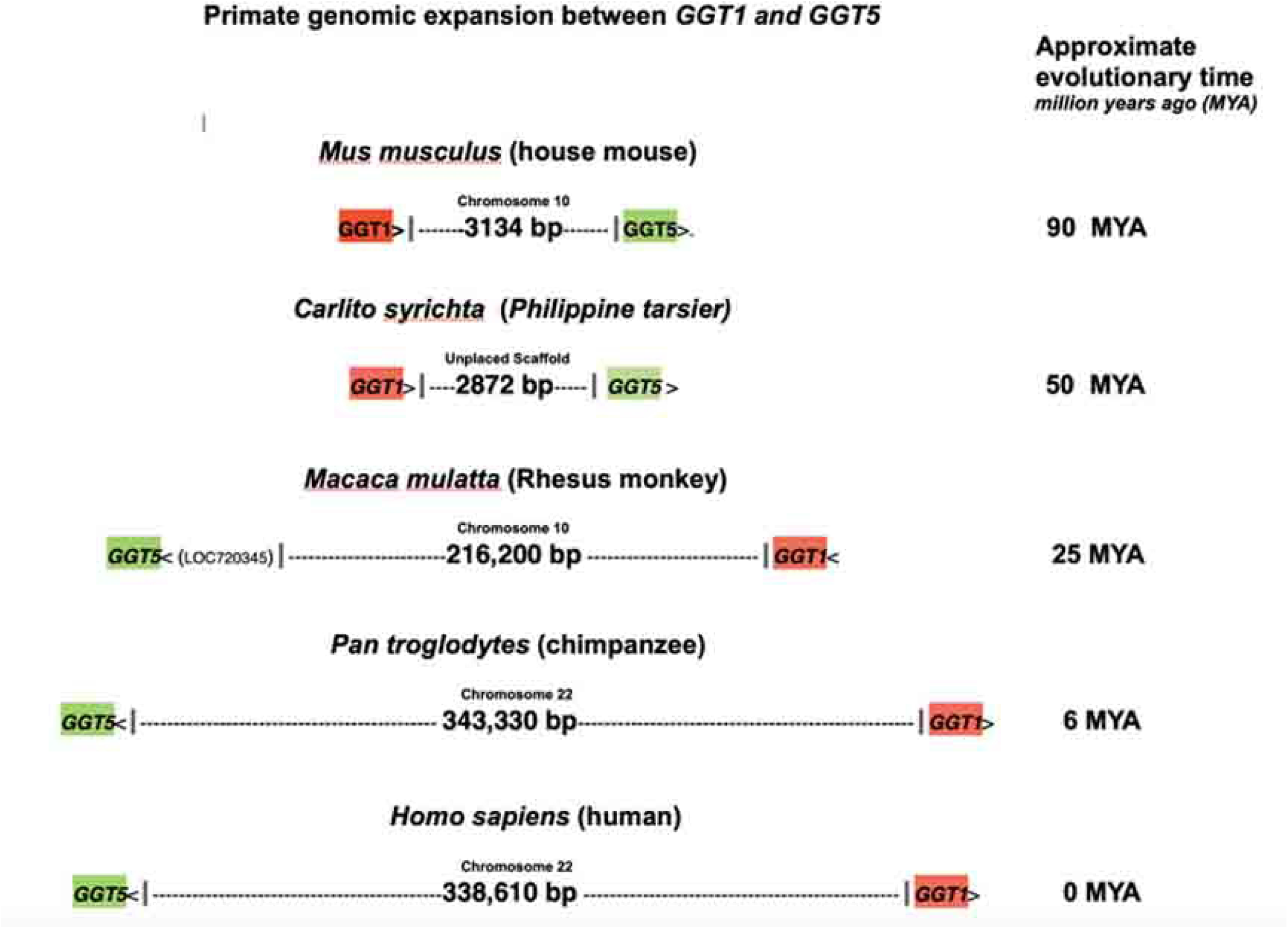
The spacer region/genomic lengths between *GGT1* and *GGT5* in various species. The house mouse and Philippine tarsier (member of ancestral primates) are in the top two drawings. The lengths of genomic regions between genes *GGT1* and *GGT5* in the higher primates are shown below. The chromosomal region is inverted in Rhesus and other primates. Chromosomal locations are also shown above the schematics. The approximate evolutionary time is on the right. Genomes of these species were analyzed from the NCBI data base (https://www.ncbi.nlm.nih.gov).

Fig. 2a depicts several diverse genes in genomic regions that follow the 3’ ends of the *GGT1*-spacer sequences, and these genes are present in different species.

**Fig. 2.**
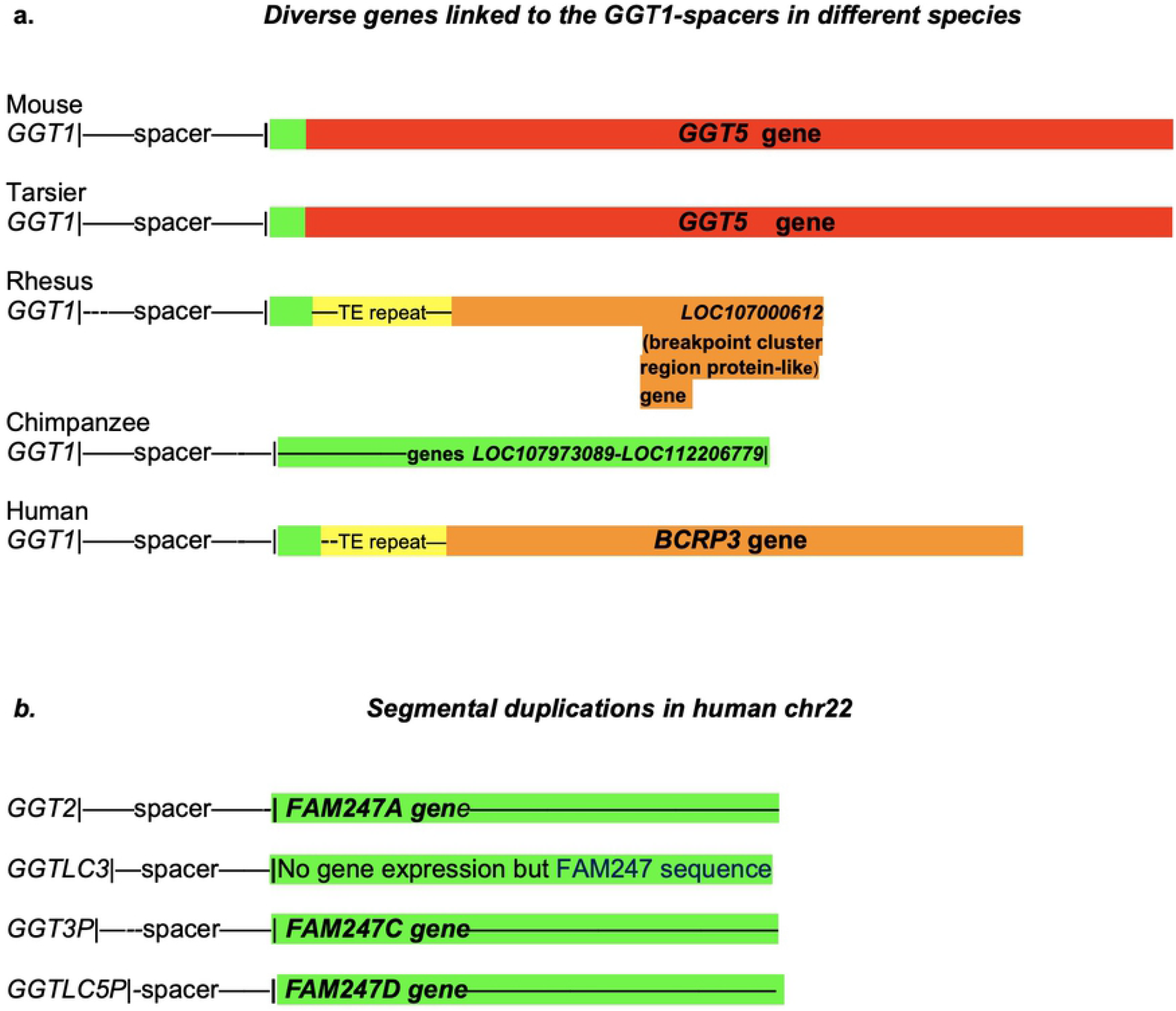
**a.** Diverse genes (*GGT5*, *LOC107973089, LOC112206779, LOC107000612, BCRP3* and FAM247) present in different species that stem from *GGT1*-spacer sequence 3’ ends. The FAM247 sequence (highlighted in green) is partially or totally present in all genes and/or sequences linked to spacers. The yellow highlighted regions represent Alu/LINE TE tandem repeat arrays. The tan areas contain sequences of the *BCR* gene. **b.** Diagrammatic representation of the spacer sequences that lead to gene sequences, The green highlighted regions represent FAM247 sequences present in different genes that start close to the 3’ ends of spacers. GGT-related genes are highlighted in re and BCR-related in tan. There are FAM247 5’ end sequences present in some spacers, e.g., mouse and tarsier but not depicted in the diagram. **b**. The *GGT*-spacer-*FAM247* gene family sequences present in different segmental duplications in human chr22 [14]. There appears to be no transcript expression from the FAM247 sequence associated with the *GGTLC3*-spacer, although the sequence has 99.4% identity with lincRNA gene *FAM2347A* and contains the entire *FAM247A* sequence [15]. GGT family genes *GGT2, GGTLC3, GGT3P, and GGTLC5P* are protein or pseudogenes that developed at duplicated loci. 25bp and 354 bp of FAM247 are not in the *GGT5* sequences of the mouse and Tarsier, respectively, and 34 and 36 base pairs of the 5’ end of FAM247 are not in the BCRP3 sequence *GGT5* genes of Rhesus and humans, respectively.

Additionally, in humans, gene duplication of the GGT-spacer motif gives rise to both *GGT*-related family genes and the *FAM247* lincRNA family genes (Fig. 2b). In terms of mechanism of initiation and growth of newly formed sequences from spacer 3’ ends, we do not know the source of the FAM247 template or the primer for DNA synthesis, or even if there is a template involved in new DNA synthesis.

The 3’ ends of the mouse and tarsier spacer sequences are defined by the start of the *GGT5* gene (Fig. 1). Because of sequence expansion in the higher primates, the *GGT5* gene locations cannot be used to define the ends of spacers. However, we have used the presence of other genes or the FAM247 sequence where there are no gene annotations to define the 3’ end of spacers in the higher primates. The 5’ half sequence of FAM247 is present in all genes/sequences that stem from spacer 3’ ends; the term FAM247 is used throughout the manuscript to denote the lincRNA *FAM247A* gene sequence or part of it. The presence of the 5’ end FAM247 sequence is helpful in estimating the 3’ ends of the spacers from species where there are no gene annotations immediately following the spacer but where the FAM247 sequence is present, e.g., in the baboon, gibbon and orangutan genomes.

Table 1 shows a high conservation of the overall bp sequence of spacer sequences between the primates, and a 56% sequence identity between the mouse and human spacers. In addition, the spacer 3’ regions display blocks of totally invariant sequences amongst the primates and the mouse (highlighted in light blue, Fig. 3). The functions of these conserved sequences are not known, but because of their invariance over evolutionary time, they may function to initiation gene sequence from the spacer 3’ end.

**Fig. 3.**
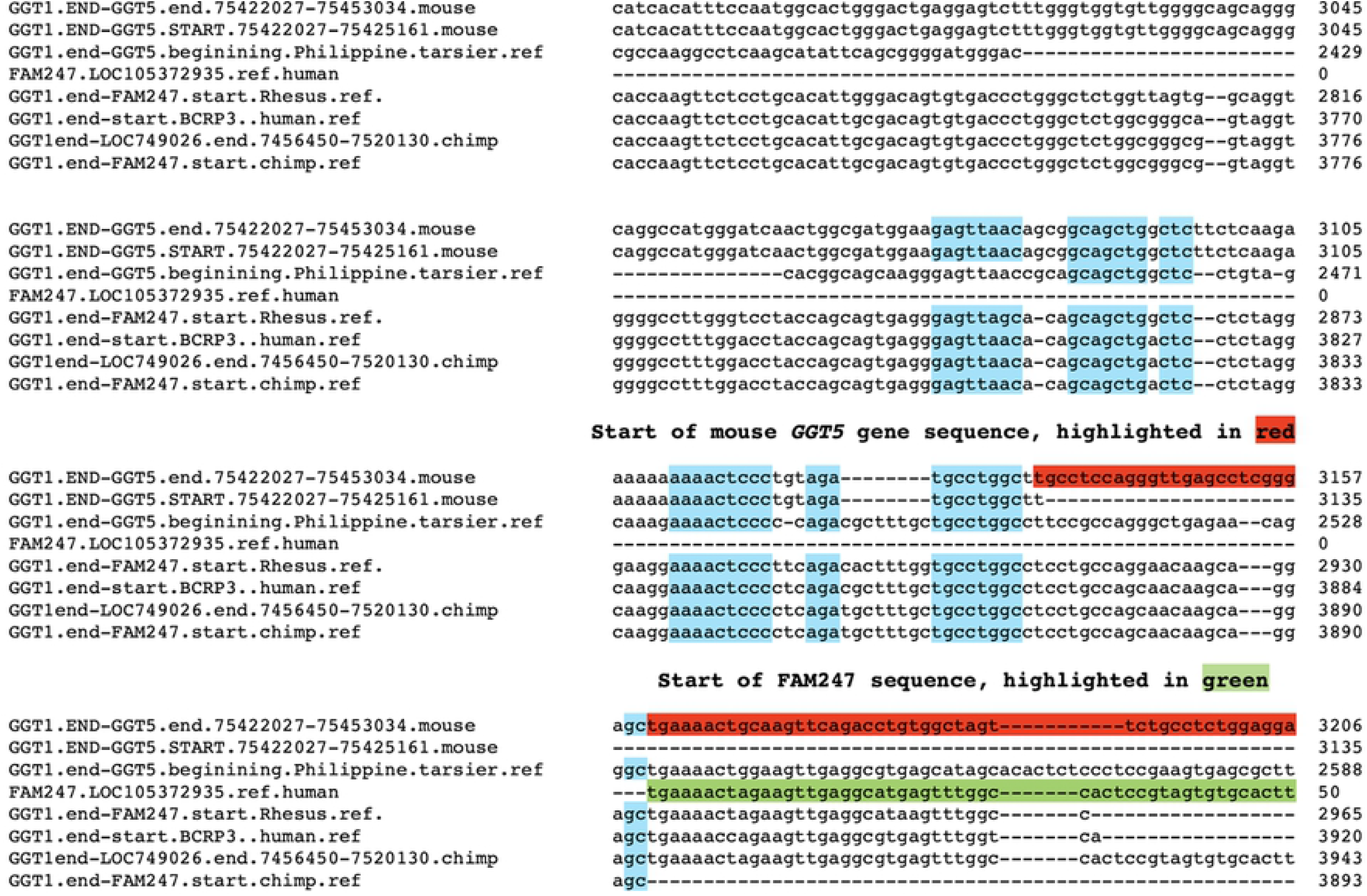
Small segment of alignment of spacer sequences showing the start of the FAM247 sequence and conserved sequences. Alignment of five spacer sequences from mouse, tarsier, Rhesus, chimpanzee and humans. Only section of the spacer 3’ terminal ends, the start of the mouse GGT5 and the start of FAM247 sequences are shown. Spacer terminal ends: mouse, at 3161 bp; tarsier, 2872 bp; Rhesus, 2933 bp; chimpanzee, 3893 bp; human, 3920 bp. Light blue highlighted, conserved sequence blocks that are conserved in all species analyzed,. Green highlighted, the start of FAM247 sequence. Red highlighted, start of *GGT5* gene sequence in the mouse genome. In the higher primates, since *GGT5* is distal to *GGT1*, the start of the *GGT5* sequence can not be used to define the 3’ ends of the spacers and the FAM247 sequence has been used. The lengths of spacer 3’ ends vary between species. However, the spacer end of the chimpanzee is shown, i.e., position 3893 bp of the chimpanzee sequence from GGT1.end-FAM247.start.chimp.ref, which ends before the FAM247 sequence begins. In humans, the *BCRP3* gene contains the FAM247 sequence but starting with position 33 of the FAM247, with positions 1-32 bp of FAM247 present in the human spacer, therefore we have defined the start of the *BCRP3* gene sequence as the human spacer 3’ end. The alignment of the complete sequences used is in S2 Fig.

**Table 1.**
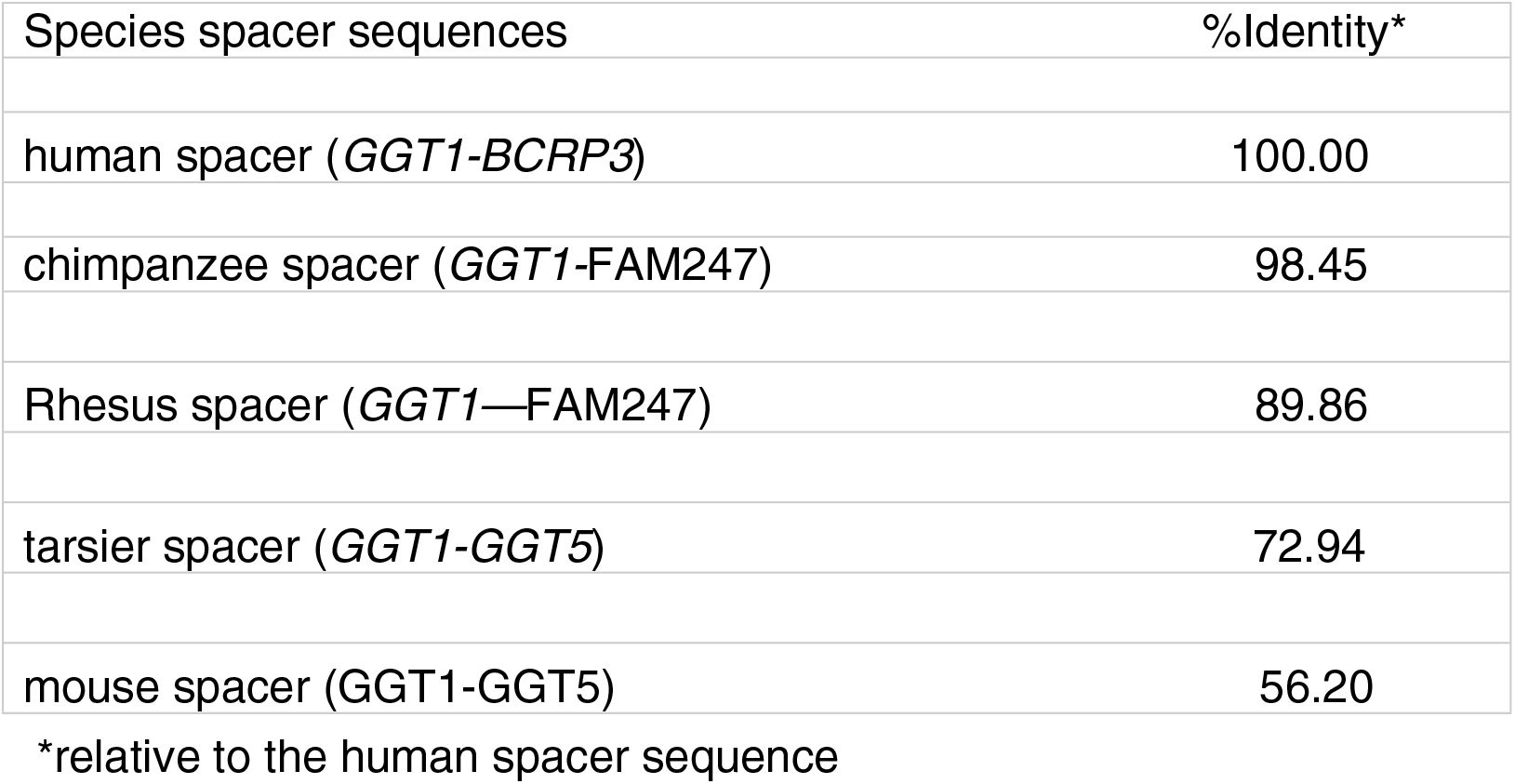
*GGT1*-associated spacer sequences and percent identity between species

There are spacer sequences between *GGT1* and *GGT5* in the zebrafish and opossum genomes, but these have significantly diverged in base pair sequence and do not display conserved sequence blocks; they have not been included in the comparisons in Table 1 or in Fig. 3.

Additional sequence conservation is within the spacer 5’ end region, and it is shown in the alignment of the 5’ end spacer gene sequences from all species considered (S1 Fig. a.).This alignment also has an added sequence, the NCBI sequence termed: GGT1 RefSeq, Homo sapiens gamma-glutamyltransferase 1 (GGT1) NG_008111.1 (website: Homo sapiens gamma-glutamyltransferase 1 (GGT1), RefSeqGene on chromosome 22). Note that the GGT1 RefSeq contains the entire *GGT1* gene sequence but also includes regions beyond the 5’ and 3’ ends of the gene, for example, the (GGT1), RefSeqGene extends 2010 bp beyond the *GGT1* gene 3’ end. The sequence alignment between spacer sequences from different species, with the extended 2010 bp sequence included, show a similarity in sequence from position 1-1451 bp of the (GGT1) RefSeqGene with sequences of the 5’ ends of the spacers, particularly with sequences from the Rhesus to human. Of significance, the distantly related prosimian primitive primate gray mouse lemur spacer shows a particularly high identity (84%) with part of the human (GGT1) RefSeqGene 3’ end sequence (S1 Fig. b); thus, a segment of the 5’ region of the spacer sequence shows a high evolutionary conservation that spans ~55 million years. The 2010 bp sequence, which follows the *GGT1* gene 3’ end, makes up a large portion of the spacer sequence in humans.

### GGT5

Of the three experimentally determined genes that are linked to spacer sequences, i.e., *GGT5, BCRP3* and the *FAM247A-D* gene family, *GGT5* is the most difficult to analyze in terms of mechanism of formation. The gene is present in the genome of zebrafish, an ancestral vertebrate species that predate the rodents and primates. However, the gene bp and protein aa sequences have significantly diverged over evolutionary time. Although there are small blocks of aa acid sequences such as _280_PPPPAGGA_287_ in the zebrafish *GGT5b* aa sequence that are totally conserved in all species analyzed, i.e., zebrafish, opossum, mouse and all primates, the overall zebrafish *GGT5b* gene aa sequence shows an identity of only 48% relative to the human sequence, showing a poor aa sequence similarity with the other *GGT5* genes. However, there is a continuum of decline of aa identity relative to the human gene aa sequence during evolution that shows a continuous sequence drift for this gene (S3 Fig). The aa sequence blocks of 100% aa identity, such as the one shown above, may be related to important functional roles of these invariant segments from the GGT5 protein. Included in S3 Fig. is the *GGT5* aa sequence of the opossum (*Monodelphis domestica*, gray short-tailed opossum), which is approximately 175 MYA in evolutionary age and thus predates the rodents. Addition of the opossum aa sequence aids in the assessment of the evolutionary changes in *GGT5* aa sequence and pattern of change and supports the continuum of evolutionary changes observed. From the *GGT5* aa sequences that have been analyzed, the data suggest that the *GGT5* genes from zebrafish to humans are evolutionarily related.

Evidence was presented to show that *GGT5* exon1 consists entirely of the FAM247 sequence in humans, primates and the mouse, but the FAM247 presence in zebrafish was uncertain [18]. Here we show evolutionary changes of *GGT5* exon1 aa sequences, with the opossum exon1 aa sequence included (Fig. 4); this helps show the trend in loss of conserved aa found with evolutionary time, but also supports the evolutionary conservation of certain aa positions, which are found to be highly biased in terms of the presence in different regions of the peptide chain (Fig. 4). There is a substantial loss of conserved aa residues in the first two thirds of the sequence, but a stability at the carboxyl terminal end of the exon1 sequence where a majority number of aa residues do not change from primates, rodents, opossum and zebrafish (Fig. 4). Thus, although the overall percent identity of *GGT5* exon1 aa sequences from zebrafish and opossum compared to that of humans is poor, the invariant aa positions of exon 1 and their highly biased locations in the peptide chain suggest an FAM247-type sequence also forms exon1 of zebrafish and opossum *GGT5* genes. Development of the zebrafish *GGT5* gene’s 5’ end sequence may have begun with the FAM247 sequence, but how the *GGT5* sequence was extended and matured to its full sequence during its birth, either in zebrafish or another early ancestor, is not known.

**Fig. 4.**
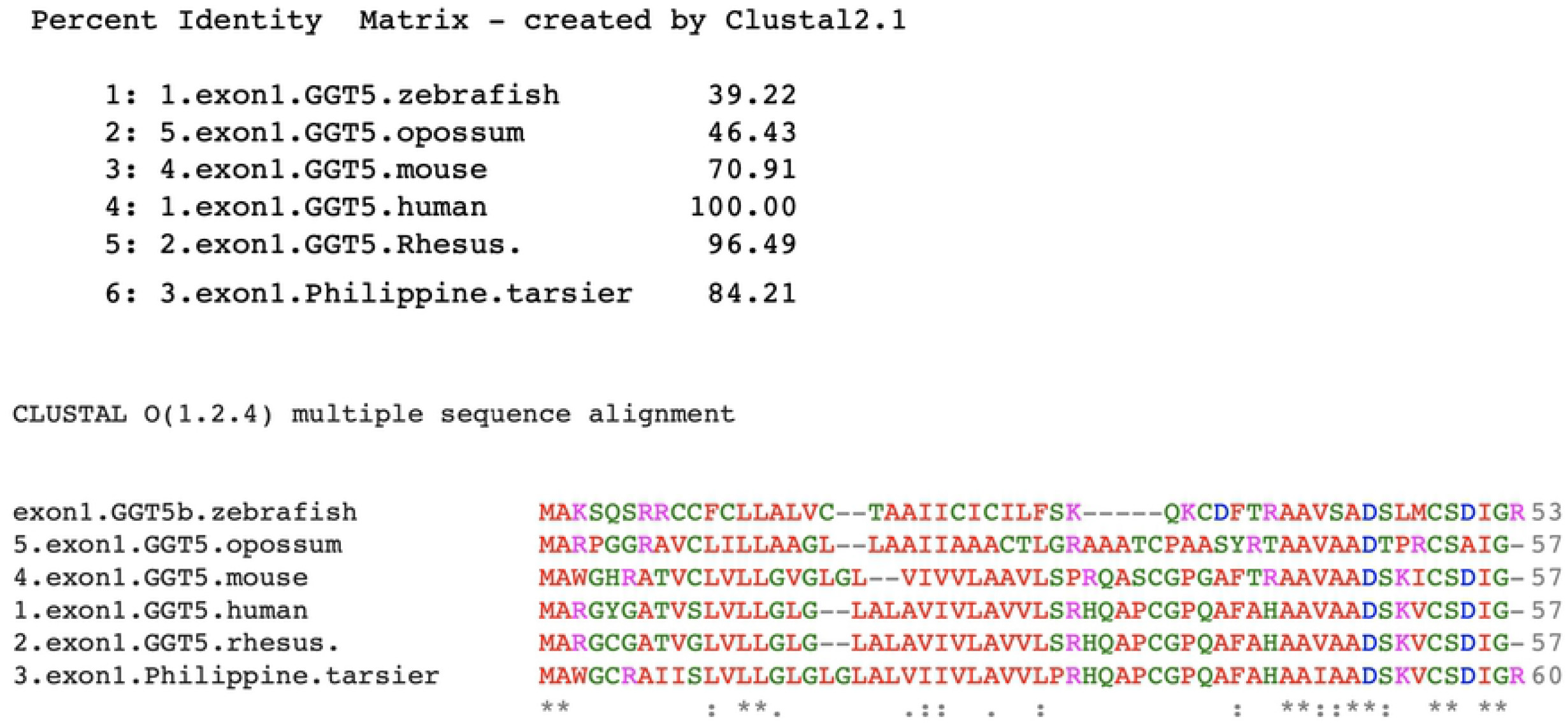
Alignment of amino acid sequences of exon 1 from GGT5 proteins from various species. Top. The percent identities between the human exon 1 and other species. Bottom. Amino acid sequences alignment showing tinvariant aa residues with *. Aligned by Clustal2.1 (https://www.ebi.ac.uk/Tools/msa/clustalo/).

### *BCRP3*: Formation of the BCRP3 sequence in Rhesus

In terms of gene expression, the human *BCRP3* gene produces one transcript that is expressed primarily in the testes (NCBI https://www.ncbi.nlm.nih.gov/gene/?term=Homo+sapiens+BCRP3) [19]. Structurally, *BCRP3* consists of approximately the 5’ half of *FAM247A* gene sequence, an Alu/LINE TE tandem repeat array, and a copy of a segment of the *BCR* (the BCR activator of RhoGEF and GTPase) gene sequence [18] (Fig. 5a). The *BCRP3* gene offers an interesting picture of how a gene sequence was created and evolved in primates over evolutionary time. Using sequence blast searches, the earliest detection of the BCRP3 sequence is in the Old World monkeys, the Rhesus monkey and baboon. In Rhesus, the BRCP3 sequence is found in chr10, linked to the *GGT1*-spacer at its 3’ end, however the BCRP3 sequence has differences; primarily, it is shorter compared to the human *BCRP3* (S4 Fig). The Rhesus BCRP3 sequence contains the 5’ half of the FAM247 sequence (with 88% identity compared to the human *BCRP3*), significant differences in the Alu/LINE TE tandem repeat array (Table 2) and a sequence segment of the *BCR* gene sequence with a significantly shorter BCR component compared to the human *BCRP3* gene (Fig. 5a).

**Fig. 5.**
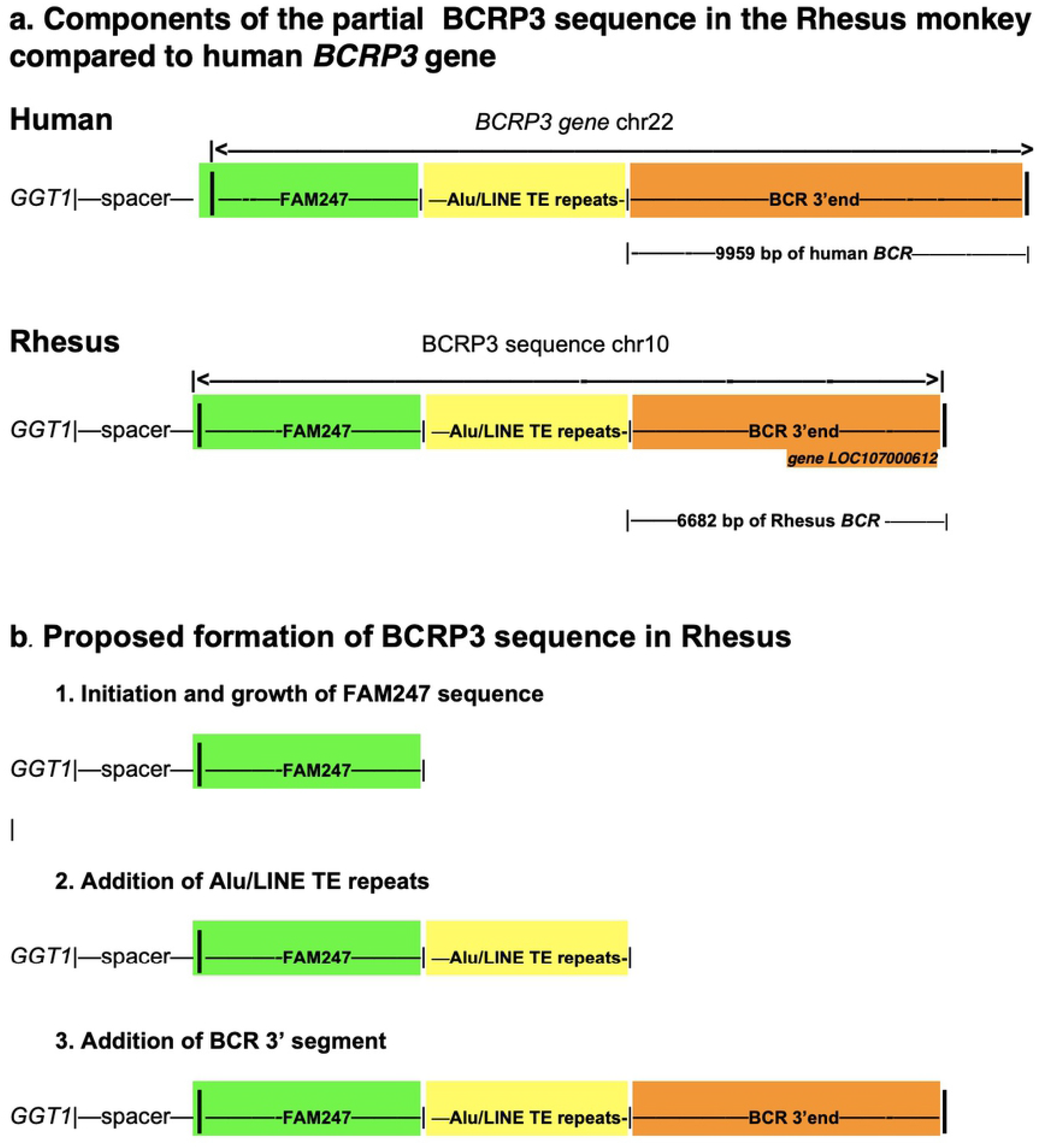
a. A schematic of the composition of the BCRP3 sequence in Rhesus compared to that of the human_*BCRP3*_gene. b. The proposed formation of the BCRP3 sequence in Rhesus, with three steps that involve initiation of synthesis and sequence growth, followed by addition of a complex of TE motifs and ending with addition of a segment of a gene from another part of the genome.

**Table 2.** Alu/LINE TE tandem arrays in BCRP3 sequences of primates*.

| Human | Rhesus | Baboon | Chimpanzee |
| --- | --- | --- | --- |
| *L1MEg* | *L1MEg* | *L1MEg* | *L1MEg* |
| MER4E1. | (AT)n | AluY | MER4E1 |
| AT)n. | AluSx | MER4E1 | (AT)n |
| (ATATACACAC)n | AluSx | (AT)n | AluSg |
| (AT)n | AluSx | AluSg | AluSx |
| AluSg | L1MEg | AluSx | AluSx1 |
| AluSx1 | AluY | AluSx1 | L1MEg |
| AluSg | AluSz6 | L1MEg | AluSx1 |
| L1MEg | L1MEg | AluSx1 | L1MEg |
| AluSx1 | AluSg4 | L1MEg | AluSg4 |
| L1MEg | (TTAT)n | AluSx | L1MEg |
| AluSg4 | AluSx | (T)n | AluSx |
| L1MEg | AluYRb3. | L1MEg | AluSx |
| AluSx | AluSx | AluSx | L1MEg |
| AluSx | AluSx | AluSx | AluSx |
| L1MEg | L1MEg | L1MEg | L1MEg |
| AluSx | AluSx | AluYRa1 | AluJb |
| L1MEg | L1MEg | L1MEg | L1MEg |
| AluJb | AluJb | AluSg | FLAM_A |
| L1MEg | L1MEg | L1MEg | MADE1 |
| FLAM_A | AluJb | L1MEg | A-rich |
| MADE1 | AluY | FLAM_C | AluY |
| (A)n | *L1M2* | A-rich | *L1M2* |
| AluY |  | *L1MEg* |  |
| *L1M2* |  |  |  |

\*data obtained by Dr. Jessica Storer using an updated RepeatMasker program

There are no genes annotated at the start of the BCRP3 sequence, but there is a gene stemming from the 3’ end of the Rhesus BCRP3 sequence, the computationally derived protein gene *LOC107000612* annotated as breakpoint cluster region protein-like (Fig. 5a); it comprises 2889 bp and is homologous to the human *BCRP3* 3’ end segment with 90% identity. Thus, a putative gene stems from the Rhesus BCRP3 sequence but the major portion of the BCRP3 sequence has no annotations, and the protein gene *LOC107000612* greatly differs from the human lincRNA *BCRP3* gene.

A model for the formation of the BCRP3 sequence in Rhesus is described below. Fig. 5b graphically shows the proposed model of BCRP3 formation.

### Model of the formation of the BCRP3 sequence in Rhesus monkey

1. Initiation and growth of the BCRP3 sequence begins at the 3’ end of the spacer with the elongation of the FAM247 sequence up to FAM247 position 5955 bp (Fig. 5b, section 1). The spacer may provide signals to initiate synthesis of the FAM247 sequence.
2. An array of contiguous Alu/LINE tandem repeats, other TEs, and AT simple repeats are added to the FAM247 sequence (Fig. 5b, section 2). Table 2 shows the TE tandem repeats. A search for a copy of a similar Alu/LINE TE tandem array in other parts of the Rhesus genome was negative. This is in contrast to the human Alu/LINE TE tandem array in the *BCRP3* gene where an almost identical TE Alu/LINE array is present in the human *IGL* locus [18]. How the TE tandem repeats were added to the growing sequence in Rhesus is not known. However, there are significant differences with TE insertions and simple repeats between the Rhesus and human TE tandem arrays (Table 2). Also, the tandem repeat in human *BCRP3:* AluSg- AluSx1- AluSg- L1MEg- AluSx1- L1MEg- AluSg4 consists of a nearly perfect tandem repeat array with no extraneous base pairs between repeating TEs, which is not the case for the Rhesus array (S5 Fig). This suggests a *de novo* formation of TE arrays with each species.
3. The Rhesus *BCR* (BCR activator of RhoGEF and GTPase) is a large gene of 133735 bp. A small section (6682 bp) of the 3’ end of *BCR* is copied and transferred to the growing Rhesus BCRP3 sequence (Fig. 5b, section 3). The segment of the *BCR* gene present in the Rhesus monkey is homologous to the human *BCRP3* gene sequence (with 82% identity) but its length is shorter than that in the human *BCRP3* gene (Fig. 5a). A copy of part the Rhesus *BCR* gene sequence may have been transferred to the growing Rhesus BCRP3 sequence linked to the Rhesus *GGT1*-spacer. There is also a partial copy of the Rhesus *BCR* sequence present in the Rhesus *IGL* locus, but it is unlikely the source of the *BCR* sequence in the Rhesus BCRP3 as the BCR fragment in the *IGL* locus is not long enough.

### BCRP3: The process of formation of the BCRP3 sequence in the baboon, gibbon and orangutan

The baboon is classified as part of the Old World monkeys and is related to the Rhesus monkey, but diverged ~2 MYA [20]. It has partially developed the BCRP3 sequence at its genomic *GGT1*-spacer locus but did not progress as far as the Rhesus in sequence development. It has the FAM247 sequence up to FAM247 position 5955 bp at 92% identity with the human *BCRP3* gene and compared with the Rhesus BCRP3 sequence at 88% and has the repeat Alu/LINE TE array (Table 2). The tandem repeats of the baboon are more similar to the human repeats than to those of the Rhesus, but missing in the baboon TE tandem array are an Alu, and MADE1 that are present in the human array at the 3’ end (Table 2). Significantly however, the baboon BCRP3 sequence differs from that of the Rhesus in that it does not have a copy of the *BCR* gene segment (S6 Fig.) and in terms of similarity of the partial sequence, it is closer to the human *BCRP3*. In addition, there are no genes annotated at the locus where the homologous partial BCRP3 sequence resides in the baboon genome. Thus, there is no apparent explanation for synthesis of the partial BCRP3 other than a failed attempt to synthesize a more complete BCRP3 type sequence or produce a sequence that can encoded a gene.

Fig. 6 summarizes the variety of sequences that stem from the 3’ ends spacer sequences in different species of the superfamily Hominoidea. The gibbons (*Nomascus leucogenys* (northern white-cheeked gibbon) are part of the family Hylobatidae, a branch of the superfamily Hominoidea (that consists of the human-like apes and humans) but are the lesser apes or small apes. Their evolutionary appearance is ~17MYA. Of major interest, at the *GGT1*-spacer locus, only part of the FAM247 sequence has formed up to FAM247 position 4467bp, which is shorter than the Rhesus FAM247 sequence at 5955 bp, but it displays a high identity with the human FAM247 sequence (95%). In addition, at this chromosomal locus, the gibbon sequence does not have a Alu/LINE TE tandem array and does not have a copy of the BCR segment of the *BCR* gene. There are no annotated genes that stem from the partial FAM247 sequence. Thus, it appears to have initiated a partial human *FAM247* gene sequence with a high identity with the human FAM247 at the gibbon *GGT1*-spacer locus, but was unsuccessful in completion of a full FAM247 sequence, the presumed end result.

**Fig. 6.**
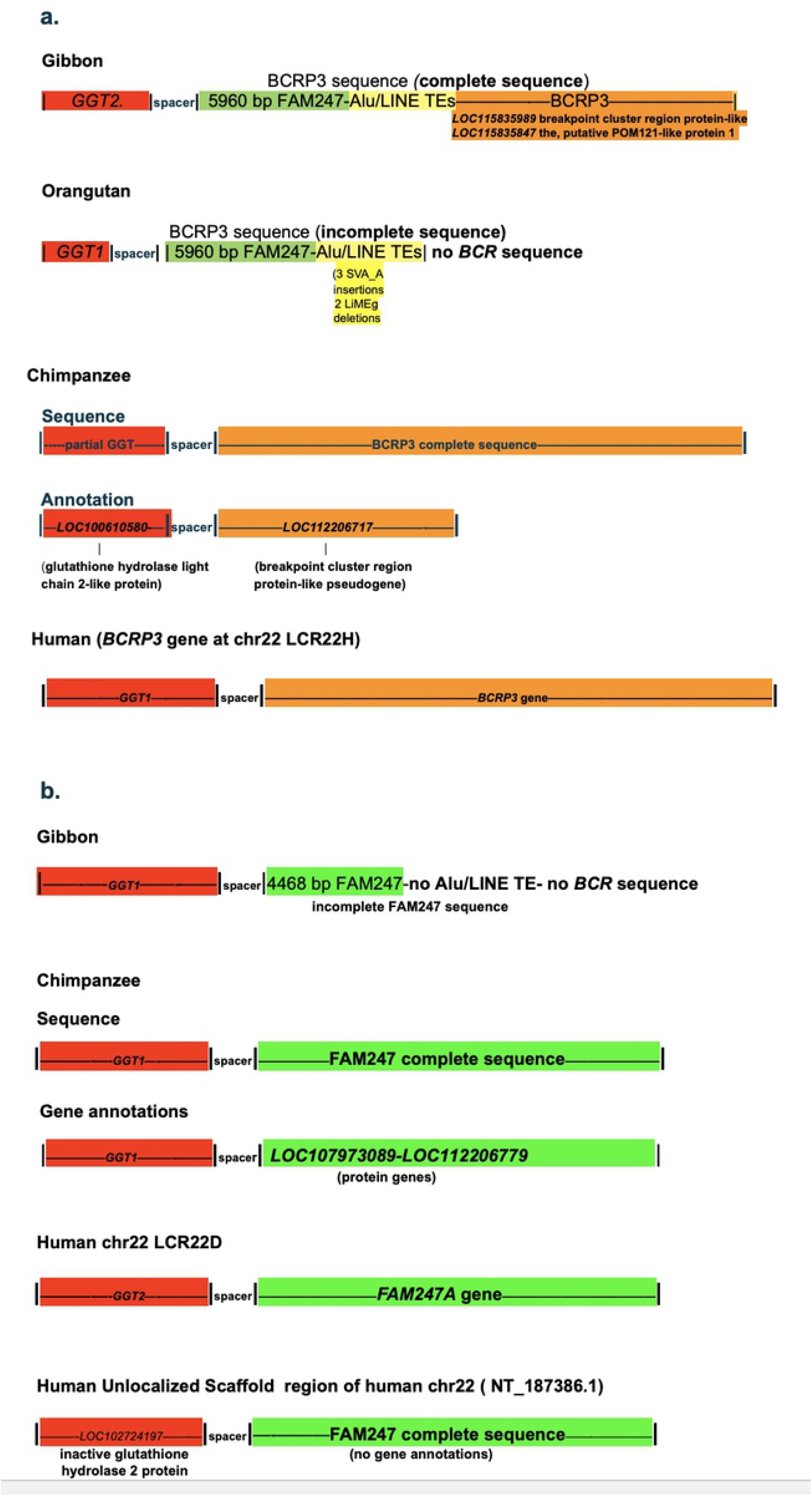
Formation of different sequences and/or genes at spacer 3’ ends sequences from species of the superfamily Hominoidea. a. BCRP3 sequence present at loci that contain a duplication of the GGT-spacer motif showing diverse genes stemming from the BCRP3 sequence. b. The presence of the FAM247 sequence at different GGT-spacer loci in gibbon, chimpanzee and human genomes.

The gibbon genome, however, has formed an almost complete BCRP3 sequence, but at another chromosomal locus, a GGT-spacer duplication locus that has the *GGT2* gene-spacer sequence (not the *GGT1*-spacer) (Fig. 6a). Although it has a base pair identity of 95% compared to the human *BCRP3*, there are several gaps in the sequence and one large additional sequence (3307bp) present in the gibbon BCRP3 sequence that is not present in the human *BCRP3* gene (S7 Fig). There are two genes annotated within part of the BCRP3 sequence, the gibbon *LOC115835989* breakpoint cluster region protein-like and *LOC115835847*, the putative POM121-like protein 1 (Fig. 6a). It appears the gibbon formed a sequence close to that of the human *BCRP3* but may have used this sequence to form two genes of its own.

The orangutans are also part of the superfamily Hominoidea and are classified with the great apes. The orangutan appeared evolutionarily about ~9 MYA. Similar to the baboon, the orangutan has formed only a part of the BCRP3 sequence at its *GGT1*-spacer sequence locus and appears to have “regressed” in capacity to mature the BCRP3 sequence compared to the Rhesus. The sequence includes the FAM247 sequence and the tandem TE repeat array, but does not have the BCR sequence that forms the 3’ end region of the Rhesus BCRP3 sequence (Fig. 6a) (S8 Fig.). In addition, the partial sequence formed by the orangutan has several small sequence repeats that may represent polymerase stuttering. It also has no putative genes that are annotated within the FAM247-Alu/line TE tandem repeat sequence. Thus, the orangutan, which is evolutionarily more advanced than the Rhesus monkey has not formed the BCRP3 sequence comparable to that of the Rhesus. Similar to the baboon, the orangutan may have come to a “dead-end” in producing a more extended or complete BCRP3 sequence.

The Alu/LINE TE repeat region of the orangutan does have major differences in the middle of the sequence compared to the human Alu/line TE tandem repeat. There are insertions of three copies of an SVA_A retrotransposon and it is missing two LiMEg elements (S9 Fig.). SVA insertions are known to affect function [21, 22]. The three SVA_A retrotransposon insertions in the orangutan sequence may be related to an inability to form a more complete BCRP3 sequence, however the baboon, which also contains no BCR sequence, does not have retrotransposon insertions in its BCRP3 sequence; thus it is unlikely the retrotransposons are the cause of the partial sequence in the orangutan.

### BCRP3: formation of the BCRP3 sequence in the chimpanzee

Similar to the gibbon, the chimpanzee genome also took an unexpected pathway with respect to BCRP3 sequence development where BCRP3 synthesis occurred at a different chromosomal locus from the *GGT1* site of synteny, at a locus that represents a duplication of the GGT-spacer motif (Fig. 6a). This locus encodes a glutathione hydrolase light chain 2-like protein gene (*LOC100610580*) and it is 130,795 bp removed from the *GGT1* chromosomal locus of synteny in the chimpanzee genome. There is a full sized BCRP3 sequence formed by the chimpanzee at this duplication site with a high identity, 98% compared with the human *BCRP3* sequence (S10 Fig.). The chimpanzee has the identical Alu/LINE TE array in its BCRP3 sequence as the human *BCRP3* gene except for differences in several subfamilies of Alus, and repeat sequences that are present in the human *BCRP3* gene and not in the chimpanzee (Table 2). The major overall differences between the chimpanzee and human BCRP3 sequences are an AluY insertion in the human sequence and an AluSx insertion in the chimpanzee; these Alus are outside of the region containing the repeat Alu/LINE TEs. Thus, the chimpanzee, together with the gibbon, formed the complete the BCRP3 sequence as opposed to the baboon or orangutan, but the chimpanzee formed a sequence that is closer to the human *BCRP3* than that of the gibbon.

The duplication locus in the chimpanzee, which has the glutathione hydrolase light chain 2-like protein (*LOC100610580*) and the BCRP3 sequence, shows a computationally predicted gene annotated as *LOC11220671*, a breakpoint cluster region protein-like pseudogene (Fig. 6a). This gene is 5323 bp in length and has a sequence that is homologous to positions 11710 bp-17033 bp of the human *BCRP3* gene; it thus has only about one quarter of the *BCRP3* gene sequence. The functions of both of the putative pseudogene *LOC112206717* and the human pseudogene *BCRP3* gene are unknown.

In humans, an inactive glutathione hydrolase 2 protein (*LOC102724197*) has been annotated in an Unlocalized Scaffold region of chr22 (NT_187386.1). The NCBI transcript table shows 14 transcripts associated with *LOC102724197*. The inactive glutathione hydrolase 2 protein gene does have a linked spacer sequence but interestingly, it also has the complete FAM247 sequence instead of the BCRP3 related sequence that is found linked to chimpanzee *LOC100610580*, the glutathione hydrolase light chain 2-like protein (Fig. 6a and 6b). An FAM247 sequence may have formed *de novo* at this scaffold region of human chr22; alternatively, the FAM247 sequence at this locus may have originated by gene duplication.

### FAM247: formation of complete sequence in chimpanzee and the gene in human

Fig. 6b shows a schematic of the FAM247 sequence that is present at the chimpanzee *GGT1* locus. This sequence has a 97% identity with the human *FAM247A* lincRNA gene sequence and contains the entire length of the *FAM247A* gene sequence (S11 Fig). The chimpanzee *GGT1* linked spacer may have served as a nucleation site to initiate FAM247 synthesis, but unlike the synthesis of the BCRP3 sequence, there was continued synthesis of FAM247 until the complete FAM247 sequence was formed. There may be signal(s) directing the addition of TE tandem repeats to an FAM247 growing sequence, but in the absence of such signal(s), there may be continued FAM247 sequence growth. However, why there is a very partial FAM247 sequence in the baboon genome is not understood. Part of the chimpanzee FAM247 sequence is annotated as two genes, *LOC107973089*, and *LOC112206779*, and both are termed uncharacterized protein genes. Thus, the computationally derived protein genes in the chimpanzee differ from the human lincRNA genes where both types of genes stem from the same DNA sequence, or part of it.

Human genome chr22 has at least four copies of the FAM247 sequence and three genes that represent the *FAM247 A,C*, and *D* lincRNA family (Fig. 2b) [15]. The evolutionary relationship of the chimpanzee FAM247 sequence to the human *FAM247* long non-coding RNA gene family is unclear. It is not known if there was *de novo* synthesis of the *FAM247* gene in humans and subsequent duplication of the linked sequence by segmental duplications (see Babcock et al for chr22 duplications [23]), or that the sequence was inherited from the chimpanzee and the FAM247 sequence developed in humans into the *FAM247* gene with minor mutations and an association with a transcriptional apparatus [24]. With respect to the GGT protein family genes and human chr222 segmental duplications, the *GGT1* gene sequence appears to have been modified to form various members of the GGT family at different chromosomal loci that consist of duplications of the GGT-spacer sequence (Fig. 6b) [14].

## Discussion

Data presented in this manuscript point to a process of initiation of *de novo* gene birth in primates that arises from an intergenic spacer sequence. This non-coding DNA sequence was evolutionarily situated between genes *GGT1* and *GGT5* in genomes of ancestral prosimian primitive primates but remained attached to the *GGT1* gene after the large primate genomic expansions. Its sequence has been conserved during primate evolution. Examples are provided that show varied sequences and diverse genes stem from the *GGT1*-spacer 3’ end, or from a duplicated spacer sequence. The data point to the spacer as a nucleation factor for initiation of new gene sequences, with the FAM247 sequence consistently serving as the starting sequence.

FAM247, whose 5’ side makes up the entire human *GGT5* exon 1 sequence, appears to also be present in the zebrafish *GGT5* exon 1, based on conserved amino acid analyses. Other data show that the 3’ end of the human FAM247, a sequence, which forms exon 11 and the 3’ UTR of the ubiquitin specific peptidase 18 (*USP18*) gene transcript [15] is present in zebrafish [18]. Thus, sections of the FAM247 sequence have been present in an early ancestor approximately 300 million years ago. The primate *GGT5* gene appears to be descendant from an early ancestor, such as zebrafish, and *GGT5* may have initially been born from a spacer sequence starting with an FAM247 type sequence in zebrafish or another ancestor. However, in terms of how the *GGT5* gene sequence was elongated and completed, this is difficult to determine with current data.

As the FAM247 sequence formed parts of genes and functional elements during evolution, it would be unusual if this sequence was an isolated example. There should be other sequences that formed parts of multiple, diverse genes and/or functional elements in different life forms during evolution, as well as the presence of other spacer-type sequences.

A model is presented to show how the long non-coding RNA gene, *BCRP3* is born in the Rhesus monkey. The process consists of the initiation of sequence growth by the spacer using the FAM247 sequence, the elongation of the FAM247 sequence, followed by addition of a complex of tandem transposable elements and ending with the transfer of a copy of the *BCR* gene segment to the newly formed sequence.

The baboon, which together with the Rhesus monkey is part of the Old World monkeys, and the orangutan that is a part of the hominoids (great apes) appear to have both come to a dead end in BCRP3 development and did not progress to the extent of the Rhesus in BCRP3 sequence maturation. Both primate species show a more limited BCRP3 sequence. In addition, the partly formed BCRP3 sequences in the baboon and orangutan genomes have no known or predict genes stemming from the partial sequences. The gibbon only formed a partial FAM247 sequence at its *GGT1*-spacer locus and with no annotated genes predicted to be encoded within the sequence. These examples suggest a trial and error process in BCRP3 and FAM247 sequence maturation for these species. The final formation of the *BCRP3* gene in humans suggests a long-term evolutionary process involving gene development. Guerzoni and McLysaght [11] previously described the *de novo* formation of primate protein genes over evolutionary time; thus, the process of long term gene development during evolution may have occured with both protein and non-coding RNA genes.

Interestingly, the chimpanzee developed both the complete BCRP3 and FAM247 sequences and with a high identity of both sequences with the human gene sequences, but these sequences were formed at different chromosomal loci from those found in humans or the sites of synteny. This leaves the unanswered question of how the *FAM247* gene sequence was formed in humans, i.e., by inheritance of the FAM247 the sequence from the chimpanzee followed by translocation of the sequence, or by *de novo* formation of the FAM247 sequence at the human locus having a GGT-spacer sequence and the RNA transcriptional apparatus to form an FAM247 RNA transcript.

The TE ALU/LINE repeats of the BCRP3 sequences have similarities to chromosomal satellite sequences, e.g., HSAT1, an element that was originally found on the Y chromosome but is also present but abundantly found on chr22 [25–27]. How the tandem TE repeats that are present in BCRP3 originated in each primate species is not known. However, McGurk and Barbash [28] have pointed out that formation of tandem arrays may begin as TE dimer insertions followed by expansion to a tandem array. Also of interest are models for the birth of genomic satellite DNA repeats [29], which may pertain to the Alu/LINE TE tandem array seen here.

We do not know the function of the Alu/LINE TE tandem arrays. With centromere and pericentromeric satellites, some play a role in heterochromatin formation in Drosophila and mammals [30]. The *BCRP3* gene is situated in a pericentromeric region of human chr 22, which may be relevant. Other and diverse roles of satellites have been outlined [31].

The evolutionary formation of the *BCRP3* sequence and gene presents a sharp contrast to the creation of the gene *linc-UR-UB*, the regulatory long non-coding RNA gene found in the human genome and believed to be involved in immune system regulation and formed by a simple transcriptional read through process [15]. This reiterates the wealth of mechanisms that life forms have used to create new genes [1–17].

Lastly, the mechanism of initiation of DNA synthesis, the DNA template for FAM247 synthesis, or if there is a template involved is a “black box”. However, Liang et al [32] studied DNA synthesis with a thermophilic restriction-endonuclease-DNA polymerase and described DNA synthesizes without a template or primer; a role in the development of genes during early evolution was hypothesized. In addition, it was shown that a hyperthermophilic archebacterial DNA polymerase can elongate palindromic and imperfect palindrome tandem repetitive DNA [33]. The FAM247 5’ end sequence begins with a small imperfect palindrome; the sequence then continues to approximately 2000 bp with sections of repetitive base pairs, and then is followed by TEs (S12 Fig.). With the *BCRP3* gene, which has part of the FAM247 sequence, the imperfect palindrome lies within the spacer sequence as the FAM247 sequence within *BCRP3* starts at bp position 33 bp of FAM247 and positions 1-32 bp are within the spacer. Can this suggest template free elongation of FAM247 synthesis? Experimental studies are needed, and the significance the of FAM247 5’ end imperfect palindrome needs to be assessed.

## Methods

### Primate species genomes

Genomic sequences of species listed were accessed using Home gene NCBI: (https://www.ncbi.nlm.nih.gov/gene) and BLAST Local Alignment (https://blast.ncbi.nlm.nih.gov/BlastAlign.cgi)

#### Species

Humans, *Homo sapiens* (NCBI:txid9606)

Chimpanzee, *Pan troglodytes* (NCBI:txid9596)

Orangutan, *Pongo abelii* (:Sumatran orangutan) (NCBI:txid9601)

Baboon, *Papio anubis* (olive baboon) (NCBI:txid9554)

Rhesus, *Macaca mulatta* (Rhesus monkey) (NCBI:txid9544)

Tarsier, *Carlito syrichta* (Philippine tarsier) (NCBI:txid1868482)

Lemur, *Microcebus murinus* (gray mouse lemur) NCBI:txid30608)

Opossum, *Monodelphis domestica* (gray short-tailed opossum) (NCBI:txid13616)

Mouse, *Mus musculus* (house mouse) NCBI:txid10090)

Zebrafish, *Danio rerio* (zebrafish) (NCBI:txid7955)

### Gene source

The NCBI/NLM data base was the source of the chromosomal locations of genes, gene annotations and gene sequences of primate and other species, Website: home gene NCBI, https://www.ncbi.nlm.nih.gov/gene

### Nucleotide and amino acid sequence alignment programs

The EMBL-EBI sequence analysis program, Clustal Omega Multiple Sequence Alignment (https://www.ebi.ac.uk/Tools/msa/clustalo/) [34] was primarily used for alignments of nucleotide and amino acid sequences as well as determining identities between sequences. The identities represent only aligned sequences and do not including gaps sequences. It should be pointed out that the percent identities can vary in comparisons of homologs with lower similarities.

Pairwise Sequence Alignment, EMBOSS Stretcher (https://www.ebi.ac.uk/Tools/psa/emboss_stretcher/) and EMBOSS Needle[34] were employed for aligning two sequences.

### Transposable elements and simple repeat analyses

RepeatMasker (http://www.repeatmasker.org/cgi-bin/WEBRepeatMasker) was employed to determine the TE Alu/LINE repeat sequences. Both search engines rm - BLAST and AB-BLAST were used. Minor differences between results from both search engines did not affect the results or conclusions. An additional related resource is the Dfam data base [35], the data base for repetitive DNA families. It should also be pointed out that there are can ambiguities in annotation of TE subfamilies [36], this was not a problem in comparing TE patterns from different species. Dr. Jessica Storer, Institute for Systems Biology, provided TE and repeat sequence data present in the BCRP3 sequence using an updated RepeatMasker program.

### RNA expression

The expression of BCRP3 expression from normal tissues from website: www.ncbi.nlm.nih.gov/gene/, human tissue-specific expression from the New Transcript table subfamilies [19].

## Availability of additional data on websites

Gene searches, gene properties, and gene transcript expression data: www.ncbi.nlm.nih.gov/gene/

HUGO Gene Nomenclature Committee: Home: https://www.genenames.org

Additional database for gene properties:

GeneCards–the human gene database: (www.genecards.org)

HGNC: (Genenames.org)

Genes and expression-site guide: https://www.ncbi.nlm.nih.gov/guide/genes-expression/

## Supporting information

S1 Fig. a. Sequence alignment of the (GGT1), RefSeqGene with the GGT1-spacer sequences from mouse and primate species. b. Alignment of sequence from the *Microcebus murinus* (gray mouse) lemur with part of the 3’ end sequence the (GGT1), RefSeqGene sequence.

S2 Fig. Alignment of complete sequences from spacers.

S3 Fig. Amino Acid sequence alignment of GGT5 from various species

S4 Fig. The alignment of the BCRP3 sequence present in the Rhesus locus that contains the sequence between *LOC106996293* and *GGT1* and human *BCRP3* gene.

S5 Fig. Alignment of the BCRP3 sequence from the baboon with the *BCRP3* gene and Rhesus BCRP3 sequences.

S6 Fig. Alignment of the gibbon sequence between GGT2 and GGT1 with the human *BCRP3* sequence.

S7 Fig. Alignment of the gibbon sequence between GGT2 and GGT1 with the human BCRP3 sequence.

Nucleotide sequence alignment of the orangutan sequence that contains the BCRP3 sequence, with the human BCRP3 gene sequence.

S8 Fig. The orangutan tandem TE repeat array showing three SVA_A insertions. Data kindly provided by Dr. Jessica Storer.

S9 Fig. Alignment of the chimpanzee sequence between genes LOC112206721-LOC112206738 (containing the GGT1-spacer duplication locus) with the human *BCRP3*.

S10 Fig. Alignment of sequence between genes *LOC112206721-LOC112206738* in chimpanzee with human *BCRP3.*

S11 Fig. Alignment of the chimpanzee sequence between *GGT1* and *LOC749026* with the *FAM247A sequence* in humans.

S12 Fig. FAM247 5’ end imperfect palindrome and repeats,

## Acknowledgements

I am grateful to Dr. Jessica Storer, Institute for Systems Biology, Seattle, Washington for providing updated RepeatMasker data for transposable element analyses.

